# Split or spread - A spatio-temporal framework for the evolution of annual *Arabis* (Brassicaceae) in Eurasia

**DOI:** 10.1101/2025.10.08.681121

**Authors:** Christiane Kiefer, Marvin Zangl, Marcus A. Koch

**Affiliations:** Heidelberg Centre for Organismal Studies (COS) Heidelberg, Biodiversity and Plant Systematics, Heidelberg Universitiy, Heidelberg, Germany; Institute for Ecology, Evolution, and Diversity, Goethe University, Frankfurt am Main, Germany

## Abstract

Life history in plants appears to be an important determining factor for the capacity to cope with major environmental change. Consequently, the evolutionary history of life form may have constrained significantly present-day biogeographic distribution patterns of phylogenetically related species, which had either to adapt or migrate during Pleistocene climatic fluctuations. In particular in Brassicaceae the molecular basis of monocarpic and polycarpic flowering behavior is well understood, and it is thought that both traits are reciprocally conversed. The Brassicaceae consist in total of c. 25% of monocarpic species. However, species-rich tribes such as Arabideae consist of only 8% monocarpic species indicating evolutionary and environmental constrain towards life history transition. Here we utilized whole plastid genome sequencing through next-generation sequencing (NGS) to investigate the spatio-temporal timing of the evolution of annual species in the tribe Arabideae (Brassicaceae). Based on these data, we conducted phylogenetic reconstructions, divergence time estimations, and plastid haplotype distribution analyses. Past and present ecological niche modeling was performed for monocarpic taxa from three different Arabideae clades, *Arabis auriculata*, *Arabis nova* subsp. *iberica*, and *Arabis montbretiana*. With the integration of these approaches, we test the hypothesis that the evolution of monocarpic species in tribe Arabideae at low frequency is associated with large-scale biogeographic patterns reflecting migration rather than adaptation or reversals.

## Introduction

Life history in plants is a continuum, ranging from short lived annuals which cycle rapidly through their vegetative and subsequent generative phase to long-lived perennials which - in the case of trees - can last for centuries. Globally, annual plants account for only 6% of vascular plant biodiversity (Poppenwimer et al 2023), but prevalence of annual species is connected to increased temperature and drought (Boyko et al. 2023). Also, in plant families such as the Brassicaceae which comprise mainly herbaceous but also some (sub)shrubby plants (Walden et al. 2020), life-history is diverse. Short-lived, monocarpic summer-annuals can complete their life cycle within a few weeks (e.g. *Arabidopsis thaliana* or *Erophgila verna*) while monocarpic winter-annuals need to be exposed to a prolonged cold period in order to flower thereby increasing the time they need to complete their life cycle. In biennials the time until completion of a single round of life cycle is even more increased as biennials behave like long-lived monocarpic winter annuals, germinating early in their first year and growing vegetatively through summer and winter, only flowering in their second year, increasing the duration of their life cycle. Finally, perennials in the Brassicaceae can be short- to long-lived and are usually polycarpic. They live over many years and go through repeated cycles of reproduction, flowering and setting seeds once every season. Perenniality, and in particular polycarpic growth, is often considered as the ancestral form of life history while annuality is regarded as derived (Hu *et al*. 2003; Grillo *et al*. 2009). In the Brassicaceae 26% of all described taxa are annuals (Hohmann et al. 2015). They are distributed unevenly across the phylogenetic lineages and annuals have evolved multiple times independently (Hohmann et al. 2015).

Eventhough the annual and perennial habit are very different, the recurrent evolution of the annual life history hints at a rather simple genetic basis of this complex trait. First studies had shown that *FLC* may be involved in regulating life history and the *FLC* orthologue *PEP1* from perennial *Arabis alpina* seemed to be responsible for maintaining vegetative growth after flowering thus enabling the plant to enter a new season of reproduction (Wang et al. 2009). Comparative studies then indicated that the differential expression of *FLC* orthologues in annual and perennial Brassicaceae from the tribe Arabideae is *cis*-mediated (Kiefer et al. 2017). However, interspecific hybrids generated in this study did not show dominance of one life history over the other and were hinting more at a more complex genetic basis - but still simple enough to enable the recurrent evolution of annuals in the Brassicaceae. Indeed, recently it was shown that although high levels of *FLC* can convert an annual into a perennial, three genes from the gene family containing also *FLC* modulate life history in a dosage-dependent manner and lead to different grades of perenniality on the life history spectrum (Zhai et al. 2024).

In the Arabideae, one of the largest tribes in the Brassicaceae, only 8% of the taxa are annual, which is considerably less than the family-wide average. Annuals have evolved at least four times independently from a perennial background (Karl and Koch 2013; Kiefer et al. 2017). Usually these annuals form species-poor groups of taxa living at lower elevation that are sister to species-rich perennial groups of taxa that typically grow at higher altitudes (Karl and Koch 2013). This pattern may also be the result of glaciation cycles which forced taxa living at higher elevation to retreat to refuge areas at lower altitude. For perennials during recolonization of habitats at higher altitudes in warmer periods subpopulations of taxa may have migrated into different mountain ranges which eventually led to the formation of new species. The annuals on the other hand remained at lower elevation providing larger areas for subsequent spreading and range expansion (Karl and Koch 2013).

The Arabideae as presently circumscribed (Kiefer et al. 2025) comprises 583 species distributed across 33 genera. Monocarpic species are found in seven out of 33 of these genera, and none of these genera with more than 3 species is dominated by monocarpic species except *Tomostima* basal to genus *Draba* (Karl and Koch 2013). The phylogeographic history of predominantly perennial groups like *Draba* (Jordon-Thaden et al. 2008) or species groups such as the *Arabis alpina* aggregate (Koch et al. 2006) or single isolated species such as *Arabis scabra* (Koch et al. 2020) has been studied in great detail. These studies revealed possible migration routes and hybridization scenarios for *Draba* on a global scale and refuge areas as well as scenarios of post-glacial colonization on a Eurasian or European scale for the latter two species. *Arabis alpina* nowadays shows a typical arctic-alpine disjunct distribution pattern which is the result of the advances and retreats of the ice shields during the Quaternary glaciations (Koch et al. 2006). As in many examples of taxa which recolonized previously glaciated areas (Hewitt et al. 2004; Varga and Schmitt, 2008), *Arabis alpina* shows a pattern of higher genetic diversity in Southern than in Northern regions (Koch et al. 2006). *Arabis scabra* for example shows another disjunct distribution which is also found among numerous elements of the Lusitanian flora that on the one hand occurs on the Iberian Peninsula and on the other hand in the northwestern Atlantic region, particularly Ireland and western Britain (Beatty and Provan, 2013, 2014; Beatty et al. 2015; Koch et al. 2020). This pattern, which lacks intermediate populations in central Europe, has been the subject of various biogeographic hypotheses e.g. focusing on postglacial migration via land connections across the Celtic Sea which existed until approximately 16,000 years ago (Lambeck 1996; Koch et al., 2020). But while these analyses all target perennial taxa, studies addressing the rare annuals within the Arabideae are largely lacking. *Arabis auriculata,* a sister lineage to the clade containing other annuals and also *Arabis alpina,* shows a broad distribution that reaches from the Iberian Peninsula and North Africa to west and center-west Asia as well as the Arabian Peninsula (https://powo.science.kew.org/taxon/urn:lsid:ipni.org:names:278028-1) and has been registered as invasive species in Belgium (Desmet et al. 2020). Despite its broad distribution and (regionally) frequent occurrence phylogeographic patterns in this taxon have not been analyzed and the species has only been included in some studies on genome evolution or chromosome number (Mandakova et al. 2020; Hoffmann et al. 2010) or comparative expression of *FLC* (Kiefer et al. 2017).

*Arabis montbretiana*, *Arabis iberica* and *Arabis kennedyae* form the annual sister group of the *Arabis alpina* aggregate (Karl et al. 2012). The group shows a disjunct distribution with *Arabis iberica* being endemic to the Iberian Peninsula (GBIF: https://doi.org/10.15468/dl.pphjbh), *Arabis kennedyae* being endemic to Cyprus and *Arabis montbretiana* (GBF: https://doi.org/10.15468/dl.7v2hnu) being native to Asia Minor and adjacent regions along the Caspian Sea and further East (by country delimitation Afghanistan, Iran, Lebanon-Syria, Turkey and Uzbekistan). The genome of *Arabis montbretiana* was sequenced as the taxon was part of the annual-perennial cross performed by Kiefer et al. 2017 and the subsequent use of the resulting NILs for analysing flowering time related traits (Madrid et al. 2021; Hyun et al. 2019) and some genome size estimated were performed (Hoffmann et al. 2010). The genome of *Arabis iberica* was also sequenced and the taxon was part of comparative studies (Kiefer et al. 2017; Madrid et al. 2021). Maybe *A. kennedyae*, with the smallest distribution range of the three closely related annuals, was studied in most detail in respect to genetic diversity, reproductive biology and population and habitat monitoring (Andreou et al. 2011). Despite these genomic studies which mainly served to pull out single genes and some insights into the biology of *A. kennedyae*, no work has been performed on annual *Arabis* species.

Inhere we aim at elaborating on our earlier findings in tribe Arabideae (Karl and Koch 2013; Karl and Koch 2014) that monocarpic species from tribe Arabideae exhibit not only significantly different trait and character combinations but also show remarkably different spatio-temporal distribution range dynamics. Therefore, in our study we focused on obtaining insights into the phylogeographic histories of annual *Arabis* lineages by means of divergence time estimates based on whole plastome data and network reconstruction based on plastidic markers. We also analysed ITS1 and ITS2 (“ITS region”) by phylogenetic reconstruction for comparison with the plastid data in order to identify hybrids. Further we have applied ecological niche modeling to add to the understanding why some taxa were able to develop continuous distribution ranges while others stayed confined to comparably smaller areas. As a taxonomic consequence of phylogenetic and biogeographic considerations we also change the taxonomic rank of *Arabis nova* ssp. *iberica* and raise it to species level.

## Materials and Methods

### Plant Material

Within this study we focused on annual lineages within the Arabideae, particularly with a focus on relatives of the perennial model species *Arabis alpina*. In total we analyzed 130 accessions of which 62 were sequenced by Illumina sequencing (either genome skimming or resequencing) and seven were sequenced in other studies also by Illumina technology (Hendriks et al. 2023) (Supplementary Table 1). The remaining 61 accessions of which the *trn*LF region and ITS region had been sequenced by Sanger sequencing had been included in previous studies (Karl and Koch 2013; Koch et al. 2016). The accessions which had been sequenced by Illumina sequencing were used for assembling full plastomes by a reference-based approach and coding sequences were used for phylogenetic reconstruction and BEAST analysis while all accessions were used for generating trees based on the *trn*LF region or the ITS region for placing all material into the same context before analysing *trn*LF haplotypes separately in a geographical context for the annual lineages.

### DNA extraction, library preparation and sequencing

Plant material was collected from herbarium vouchers (Supplementary Table 1) and DNA was extracted using the Invisorb Spin Plant Mini Kit (Invisorb) according to the manufacturer‘s protocol. Sequencing libraries were prepared using the Illumina TruSeq Kit (Illumina, San Diego, CA, USA) at the Deep Sequencing Core Facility (Heidelberg University) and were subsequently sequenced on an Illumina NextSeq 550 (150bp paired end, library insert size 400bp). For HEID502973, HEID503390, HEID505446 and HEID8006710, four library preparations and sequencings (also 150 bp paired end) were performed by Novogene Co. Ltd. (UK). In summary 800,000 and 91 million sequencing reads were obtained per sample, and details are given with Supplementary Table 1.

### Reference based assembly of plastid genomes

Raw reads were trimmed using Trimmomatic v0.40-rc1 (Bolger et al. 2014) to remove sequencing adapters using standard settings. Reads with a length <50 bp were discarded. For reference-based assemblies of the plastid genome trimmed reads were mapped to a plastid reference (*Arabis alpina*, NC_023367; Melodelima and Lobréaux, 2013) using BWA v0.7.17-r1188 and the mem algorithm (Li and Durbin, 2009; Li 2013). The resulting sam file was converted to bam format by samtools v1.16.1-35 (Li et al., 2009), unmapped or ambiguously mapped reads were removed by dropping reads with a mapping quality <1 and the bam file was sorted by samtools v1.16.1-35 (Danecek et al. 2021). Duplicate reads were identified and removed by using picard-tools v2.18.25 (Broad Institute) and its ‘MarkDuplicates’ function. Variant calling was performed using the tool ‘HaplotypeCaller’ of GATK v4.2.4.1 (McKenna et al., 2010). Sample ploidy [--sample-ploidy] was set to 1 (haploid). The GATK v3.8-1-0 tool ‘FastaAlternateReferenceMaker’ was used for generating new sequences based on the respective references and the detected variants.

In a last step regions with a coverage < 5 and a mapping quality < 30 were identified using the GATK v3.8-1-0 tool ‘CallableLoci’ and masked by inserting ‚N‘ by our inhouse script masker (Kiefer et al. 2024). The masked sequences were annotated by alignment using the annotation of the respective reference sequences and the inhouse script cpanno (Kiefer et al. 2024). Annotated plastid genomes were saved in GenBank format.

In addition to the samples sequenced within this project the plastid genome of Pseudoturritis turrita (genebank accession number ERS2461510) was also assembled and treated in the same way as the other accessions in order to use it as outgroup in phylogenetic analysis.

### *De novo* assemblies of the Internally Transcribed Spacer (ITS) region

For the ITS region *de novo* assemblies of the trimmed sequencing reads were prepared using SPADES version 3.13.1 and standard settings. The resulting assemblies were used to construct blast libraries using the blast suite version 2.9.0+ (Camacho et al. 2009). The libraries were searched for contigs containing the ITS1 and ITS2 as well as the 5.8s rRNA region by blastn from the blast suite version 2.9.0+ (Camacho et al. 2009) using the *Arabis alpina* ITS sequence with the genbank accession number AJ232923 as query sequence. This approach generated virtual ITS sequences and has been successfully tested and applied throughout the entire Brassicaceae family (Kiefer et al. 2025).

### Phylogenetic reconstruction based on plastome, *ITS* and the *trn*LF region (*trn*L gene, intron and *trn*L-F intergenic spacer) data

For phylogenetic analysis of the plastid genomes protein coding sequences as well as tRNA and rRNA encoding sequences were extracted from the newly generated plastomes. To do so the genebank files containing the annotation information of the plastid genomes were first converted to gff format using seqret v6.6.0.0 (Rice et al., 2000). Then the annotation table was converted to bed format by gff2bed from BEDOPS v2.4.37 (Neph et al., 2012) and then filtered for intervals representing coding sequences or intervals corresponding to tRNA or rRNA encoding sequences (77 protein coding genes, 29 tRNA encoding genes and 4 rRNA encoding genes). Subsequently the intervals defined in the bedfile were extracted from the newly generated plastid sequences using BEDTools v2.30.0 (Quinlan 2014) with the ‘getfasta’ option. For genes with multiple exons, exons were fused after extraction to form continuous coding sequences (rps12, tRNA-Ala-UGC, tRNA-Ile-GAU, tRNA-Leu-UAA, tRNA-Val-UAC). Sequences were aligned for each extracted interval individually for all accessions including the outgroup by MAFFT v7.511 (Katoh and Standley, 2013) and checked in Aliview v1.28 (Larsson, 2014). Alignments were then concatenated (Supplementary File 1) using catfasta2phyml v1.2.0 (https://github.com/nylander/catfasta2phyml).

ModelFinder in iqtree2 (Minh et al. 2020; Kalyaanamoorthy et al. 2017) was used for determining the best molecular evolutionary models and partitions and iqtree2 was used for phylogenetic reconstruction (Minh et al. 2020). 500 bootstrap replicates were generated.

FigTree v1.4.4 (Rambaut, 2018) was used for visualizing the best tree and figures were edited using Adobe Illustrator CS4 v14.0.0 (Adobe Systems Inc., 2008).

For phylogenetic reconstruction based on the *ITS* region the contigs containing the best blast hits were extracted from the de novo assemblies and aligned using MAFFT v7.511 (Katoh and Standley, 2013; alignment in Supplementary File 3). The alignment ends were clipped according to the reference. Further, the alignment was supplemented with sequences downloaded from GenBank representing additional accessions of *Arabis montbretiana*, *Arabs iberica*, *Arabis nova*, *Arabis auriculata* and *Arabis verna* for obtaining a nuclear perspective on the phylogenetic placement of mainly *Arabis iberica* but also of the other species, as the maternal plastidic perspective and the nuclear signal that also includes the paternal perspective, may differ from eachother. Phylogenetic reconstruction followed the same approach as desccribed above.

For the phylogenetic placement of *Arabis nova* ssp. *nova* additional trees based on the *trn*LF region (*trn*L gene, intron and *trn*L-F intergenic spacer) were calculated. For this analysis the respective regions were cut from an alignment containing the complete reference based assembled plastid genomes using Aliview v1.28 (Larsson, 2014) and seven *Arabis nova* ssp. *nova* accessions were added to the alignments (A18MO2 (no ITS), A32M, A33M, A34M and MA024 (Karl 2009; Karl et al. 2012), RK125 (Karl and Koch, 2014) and RK127 (Karl and Koch 2013). Accession details are given in Supplementary Table 1. Sequences were realigned along with the newly added sequences using again mafft v7.511 (Katoh and Standley, 2013; alignment in Supplementary File 4). Phylogenetic trees were reconstructed using IQtree v1.6.12 (Nguyen et al., 2015). An extended model selection with ModelFinder (Kalyaanamoorthy et al., 2017) was performed within iQtree and bootstrapping was performed by using the ultrafast bootstrapping algorithm as implemented in iQtree (Hoang et al., 2018) with 1000 replicates. Further, an SH-like approximate likelihood ratio test (SH-aLRT) (Guindon et al., 2010) was run with 1000 replicates for obtaining further information on branch support.

### Divergence time estimates

Divergence time estimates for the plastid tree were performed with BEAST2 v2.6.7 (Bouckaert et al., 2019). The combined alignment and partition written according to the results from ModelFinder were used to create a NEXUS file with 8 partitions. The input file was generated by BEAUti2 v2.6.7 from the BEAST2 package. The birth-death-model (Kendall, 1949; Yule, 1997) was chosen, such as a relaxed-clock-model (Drummond et al., 2006). Trees and clocks were linked. Four independent runs were started with a chain length of 10,000,000 each. Secondary calibration was done, using two priors that were taken from (Walden et al., 2020): 1) A. alpina – D. nemorosa 2) Aubrieta verna – Aubrieta vulcanica (Suplementary Table 2). The evolutionary models for every partition were set manually (Supplementary File 2) and the tree rooted with the outgroup by setting all taxa except for Pseudoturritis turrita as monophylietic group. For partitions 1 and 7 the model (K3Pu) was not available in BEAST2, therefore the model was approximated starting from the GTR model. Resulting trace files (from 6 runs) were inspected in Tracer v1.7.2 (Rambaut et al. 2018). All trees were combined with LogCombiner v2.6.7 (Bouckaert et al., 2019). A burnin of 10% was set and TreeAnnotator v2.6.7 (Bouckaert et al., 2019) was used to obtain one target tree as a Maximum Clade Credibility (MCC) tree. Node heights were set to mean heights. The resulting tree with divergent time estimates plotted as bars was visualized in FigTree v1.4.4 (Rambaut, 2018) and edited for better readability in Affinity Publisher 2 (Serif (Europe) Ltd.).

### TCS network analysis and spatial visualization

Two subsets were extracted from the *trn*LF region alignments which had been subjected to phylogenetic reconstruction: one comprised only A. auriculata accessions while the other one contained the accessions representing *A. montbretiana*, *Arabis kennedyae* and *Arabis iberica*. These alignments were used for haolotype netweork reconstruction using TCS1.21 (Clement et al. 2000) setting gaps to missing. The resulting networks were re-drawn using the program Draw (LibreOffice 6.4.7.2) and circle sizes were adjusted for number of samples included in a haplotype. Haplotype distribution was plotted on maps using QGIS. Maps were finalized using the program Affinity Publisher (Serif, Europe, Ltd.).

### Species distribution modeling

For obtaining an insight into past and present distribution areas for *A. montbretiana*, *A. iberica, A. auriculata* and *A. nova* ssp. *nova* we performed species distribution modeling. In a first step geo-referenced point data were downloaded from www.gbif.org for the four taxa (*A. montbretiana*: https://doi.org/10.15468/dl.7v2hnu, *A. iberica*: https://doi.org/10.15468/dl.pphjbh, *A. auriculata* https://doi.org/10.15468/dl.nfj4dx, *A. nova* ssp. *nova*: https://doi.org/10.15468/dl.w8y8t6). Data were filtered using a custom script in R following the recommendations in https://data-blog.gbif.org/post/gbif-filtering-guide/ and plotted on a map using QGIS (http://qgis.org). Two datasets (recent and LGM) containing 19 bioclioclimatic variables each were obtained from worldclim.org. The bioclim raster layers were clipped according to the distribution range of *A. auriculata* as determined by the gbif data. The clipped rasters were exported as ascii files and used as input in MaxEnt 3.4.3 (Phillips et al. 2006; https://biodiversityinformatics.amnh.org/open_source/maxent/) together with the filtered occurrence data for the four species (in separate runs). Auto features was unclicked and in addition to the default options threshold features were selected. Response curves were set to be created, a jackknife analysis was chosen for measuring variable importance and the output format was set to logistic. A random seed was chosen and 10 replicates were run for cross validation.

After this initial analysis the clipped bioclim ascii files were imported into R and Pearson’s correlation coefficient was calculated using a custom script. Of any variable with a correlation coefficient > 0.75 only the one was kept which was more informative according to the first MaxEnt analysis. Subsequently the analysis was repeated for all for taxa with the remaining bioclim variables and projections were run for the LGM climate model (Pleistocene; ∼22,000 years ago). The resulting species distribution models were visualized in QGIS.

## Results

### Integration of phylogenetic reconstruction, BEAST analysis and geographic haplotype distribution

Our aim in this analysis was to integrate phylogenetic reconstruction, divergence time estimates and geographic distribution of haplotypes for obtaining an insight into the colonization of Eurasia by the annuals *Arabis auriculata*, *Arabis iberica* and *Arabis montbretiana*. Phylogenetic reconstruction based on gene sequences extracted from full plastomes comprised 69 accessions representing most lineages of the Arabideae for which full plastomes were available (Supplementary Table 1), and *Pseudoturritis turrita* (ERS2461510) served as outgroup. Phylogenetic reconstruction by maximum likelihood yielded a well resolved tree which was congruent to previously published data (Supplementary Figure 1). Seven of the eight clades described in Karl and Koch 2013 were recovered (Supplementary Figure 1) which were labeled accordingly. Bootstrap support was high (90 - 100%) across the entire tree with the exception of one node within the *A. auriculata* clade.

The BEAST analysis was also in agreement with previous analyses (Figure 1) (Walden et al. 2020). Root height and thus crown age of the tribe Arabideae was estimated to be 16.9 my. Most of the main clades originated in the Miocene or early Pliocene between 15.4 and ∼5.5 mya. Accordingly, diversification of the tribe Arabideae into the various genera as actually defined taxonomically occurred well before the onset of the Quarternary Pleistocenic glaciation and deglaciation cycles and are exclusively placed into the much warmer phases of the Miocene or in one case (*A. alpina* / *A. nordmanniana*) in the Pliocene, while the *A. alpina* aggregate and its annual sister group separated in the Pliocene.

**Figure 1.**
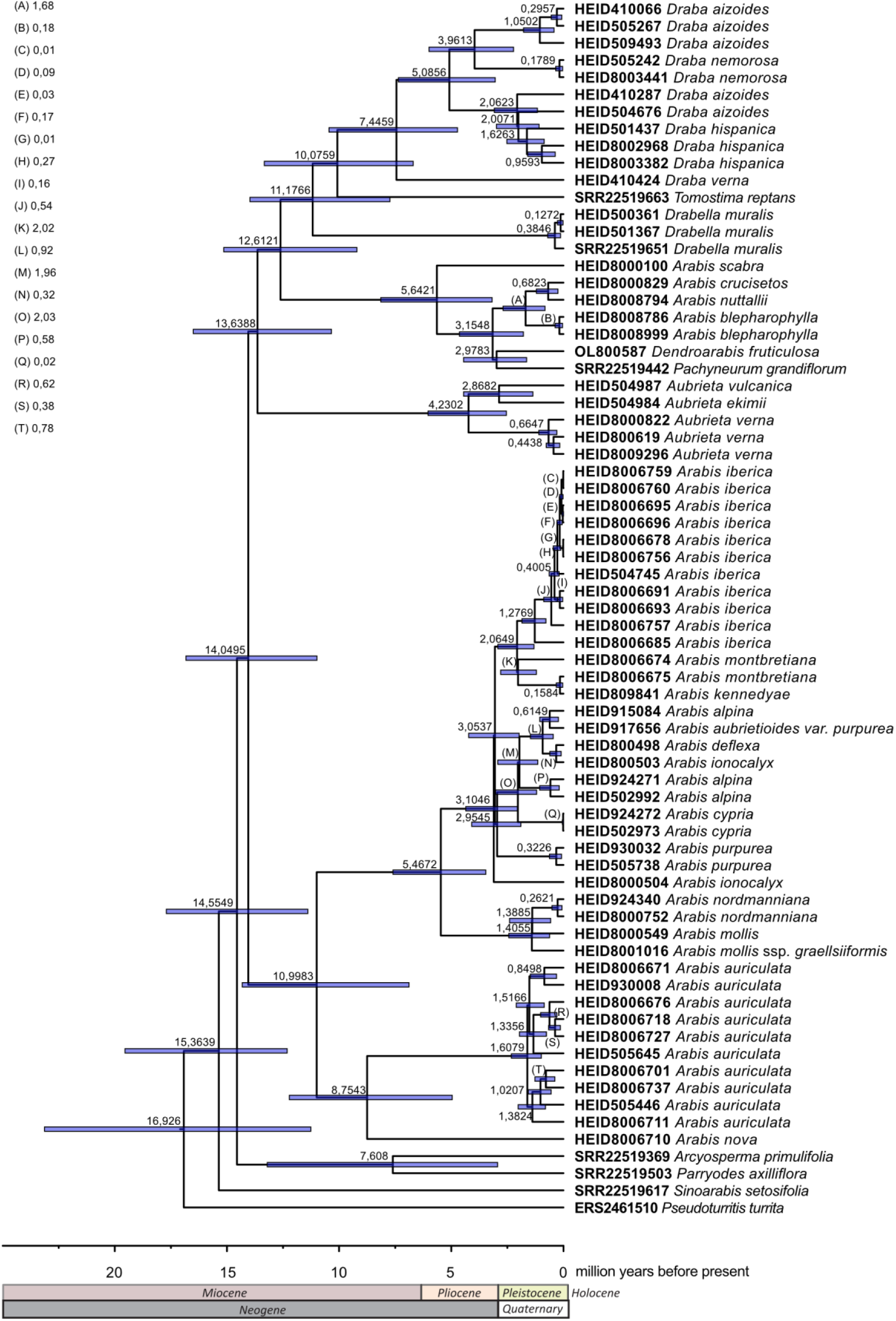
BEAST analysis using coding sequences from whole plastid genomes comprising all main lineages of the Arabideae. Four BEAST runs with 10 mio generations each were combined (every 10.000^th^ generation sampled).

Nearly all splits between accessions of the same species happened within the Pleistocene. Interestingly, while most lineages had rather short branches, the branch leading to *A. auriculata* was very long, indicating extinction of earlier lineages of the same or a closely related taxon.

For analysing the geographical distribution of haplotypes for *A. auriculata* on the one hand and the monophyletic group comprising *A. iberica*, *A. kennedyae* and *A. montbretiana* on the other hand haplotype networks were generated from alignments including only the *trn*LF region sequences of the respective taxa and plotted on maps (Figure 2A and C).

**Figure 2.**
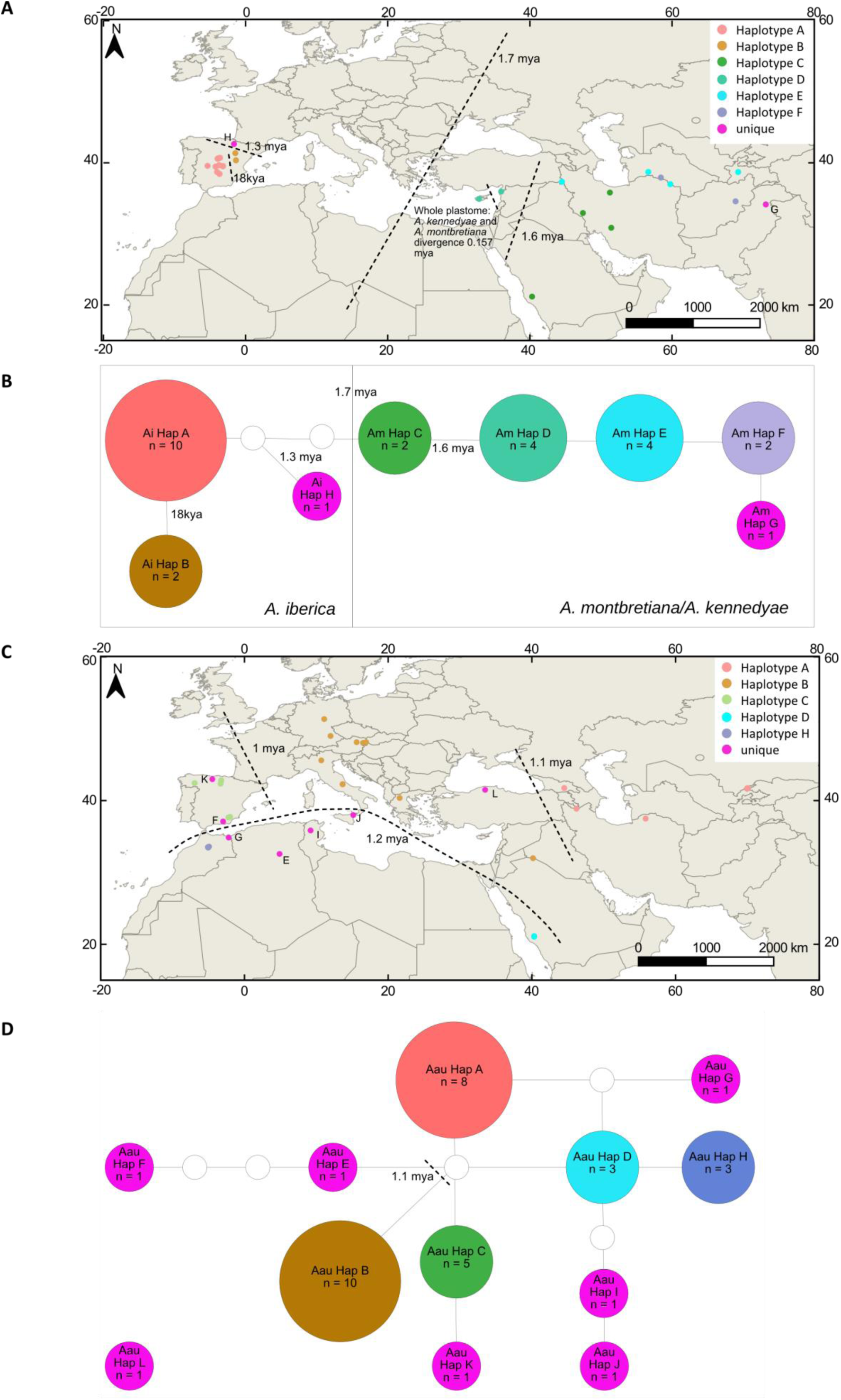
Haplotype distribution (*trn*LF region) and haplotype networks for two annual lineages. (A) Haplotype distribution for the lineage comprising *Arabis iberica*, *Arabis montbretiana* and *Arabis kennedyae*. *A. iberica* and *A. montbretiana* split ∼1.7 mya while *A. kennedyae* is much younger (split 157 kya in the whole plastome data; inhere the taxon shares a haplotype with *A. montbretiana*) (B) Haplotype network showing the relatedness of the haplotypes in (A); (C) Haplotype distribution for the A. auriculata lineage; divergence times corresponding to the splits of haplotypes indicate a rapid migration from East to West (D) Haplotype network representing the relationships of the haplotype shown in (C) Dashed lines in (A) and (C) indicate divergence times derived from the BEAST analysis. Dates in (B) and (D) are also derived from the BEAST analysis.

For *A. montbretiana* the haplotype network shows a linear series of connected haplotypes which are all separated by one mutation (Figure 2B). “Rooting” the haplotype network between *A. montbretiana/kennedyae* and *Arabis iberica* according the phylogenetic tree and overlaying it with the BEAST analysis indicates that *Arabis montbretiana* probably evolved in the Mediterranean 1.7 mya where it split into the actual *A. montbretiana* lineage and *A. kennedyae* as a local endemic on Cyprus. From the Turkish mainland it migrated further East where haplotypes Am Hap D and E which occurr in eastern Turkey, Turkmenistan and Tadjikistan evolved 1.6 mya (Figure 2A and B). The haplotypes derived from Am Hap E are found even further East so there seems to have been a linear migration towards the East.

For *A. iberica* three haplotypes were identified, one frequent one, Ai Hap A (n=10) and two less frequent ones, Ai Hap B (n=2) and Ai Hap H (n=1) (Figure 2B). There was a weak geographical pattern with Ai Hap A occurring mostly in Central Spain while the two less frequent haplotypes were found in Eastern Spain (Figure 2A). Au Hap H split from Ai Hap A and B about 1.3 mya (Fig. 1 and Fig 2A) while Ai Hap A and Ai Hap B split about 18,000 years ago. The BEAST analysis and the phylogenetic reconstruction indicate that Ai Hap B actually evolved from Ai Hap A. Therefore, we conclude that the ancestor of *Arabis iberica* arrived on the Iberian Peninsula at some point between 1.7 mya (stem group age) and 1.3 mya (crown group age) upon which the population split into a northern group (Ai Hap H) and a group in Central Spain which split again after the maximum of the last glaciation 18,000 years ago).

For *Arabis auriculata* two main haplotypes, Aau Hap A and Aau Hap B, were identified in the network analysis (Figure 2D). Aau Hap A mainly occurs in Asia while Aau Hap B is found in central and southern Europe and also on the Arabian Peninsula (Figure 2C). Aau Hap C, which is also frequent, is confined to the Iberian Peninsula. Aau Hap D is a haplotype of lower frequency derived from Aau Hap A or B through missing haplotypes that occurs on the Balkans and the Arabian Peninsula while Aau Hap H derived from Aau Hap D occurs in Northern Africa. In total six unique haplotypes were found which all occur on the Iberian Peninsula and in Northern Africa and Sicily. The phylogenetic analysis based on full plastomes suggests that haplotypes Aau Hap A, Aau Hap B, Aau Hap C and Aau Hap K form a group while Aau Hap G, Aau Hap H, Aau Hap J and Aau Hap F form the second group. According to the Beast analysis these groups split about 1.2 mya. Three haplotypes from the first group are frequent and occur in large geographical regions ranging from Central Asia through the Northern Mediterranean to Central Europe and the Iberian Peninsula (Figure 2C). The Central Asian group (Aau Hap A) split from the group that would reach Europe around 1.1 mya. The Iberian group split from the Northern Mediterranean/Iberian group about 1 mya.

The second group which occurs in North Africa as well as on the Arabian Peninsula is defined by diverse haplotypes of low frequency (Figure 2C). In the phylogenetic reconstruction the deepest split is found between haplotype F and the rest of the North African group. Haplotype F is a singleton and occurs in Morocco. However, haplotype Aau Hap D was not represented in the phylogenetic analysis. The network analysis places it near a split from A/C. This would in agreement with a possible origin of *Arabis auriculata* in Central Asia from where it migrated to the Arabian Peninsula and from there along the North Africn coast towards the West on the one hand while on the other hand it migrated via Turkey into the Mediterranean and Central Europe and from there on to the Iberian Peninsula.

### Niches occupied by the annuals in this study are all different

For determining the current and past (LGM) possible distribution of the annual species included in this study we performed niche modelling. Based on recent bioclimatic variables *Arabis auriculata* showed the broadest possible distribution which essentially covered entire central and southern Europe, North Africa and reached into Central Asia (Figure 3A). The most probable distribution modelled for *Arabis montbretiana* based on recent bioclimatic data covered a large part of Central Asia (Figure 3 C) while the most probable modelled distribution range of *Arabis iberica* was confined to the Iberian Peninsula, North Africa and some small regions in Italy, Greece and Turkey (Figure 3E). Projecting the current species distribution model on bioclimatic variables reflecting climatic conditions around the last glacial maximum (∼22,000 years ago) revealed a very strong decrease in possible suitable distribution ranges. For *Arabis auriculata* there was a low probability for suitable niches in today’s Southern Great Britain, an area connected to the European mainland during the Quaternary ice ages (Figure 3B). The possible distribution range for *A. montbretiana* did now show great changes, indeed rather an increase into areas further West. However, there was a small decrease in cloglog values which can be interpreted as the probability of a plant occurring in a certain region. The available niches for *Arabis iberica* decreased dramatically around the last glacial maximum and were essentially non-existing. The biggest probability was indeed in Turkey or on the Balkans where at least nowadays the taxon does not occur and to a much lesser extent on the Iberian Peninsula in the Cantabrian region, a known refuge area during the LGM (Magri et al. 2006).

**Figure 3.**
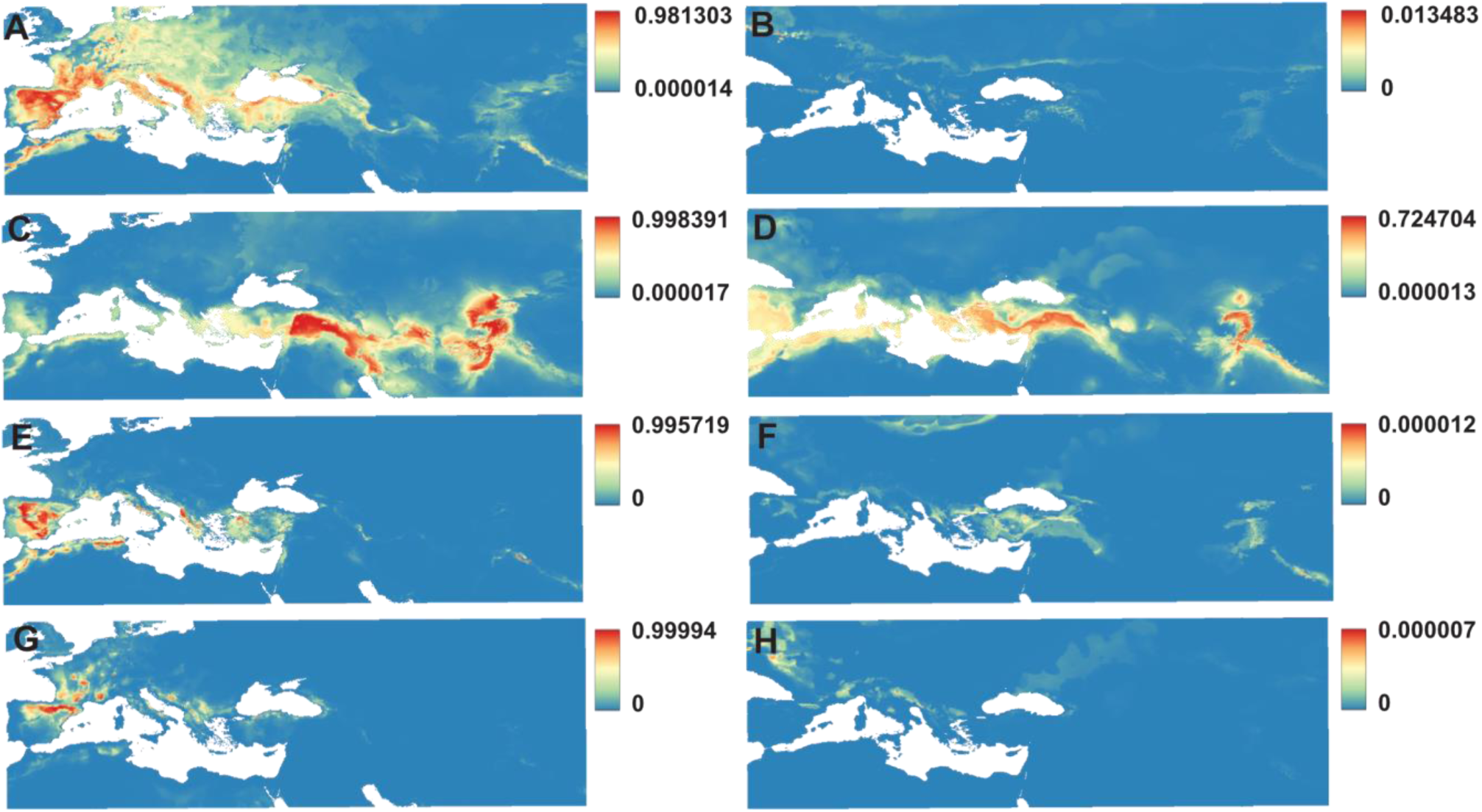
Bioclimatic Niche Modelling of four annuals. Ecological niche modelling was performed in MaxEnt (Phillips et al. 2006) using occurrences downloaded from GBIF. Modelling was performed for recent bioclimatic data (A, C, E, G) as well as for bioclim data representing the LGM (B, D, F, H). Despite their close relationship *Arabis iberica* and *Arabis montbretiana* showed very different niche occurrence patterns. Notably also *A. iberica* and *A. nova* occupy different niches which – besides the phylogenetic position – supports the description of *Arabis iberica* as an own species. (A) Modelled current distribution of suitable niches for *Arabis auriculata* and (B) distribution of suitable niches during the LGM indicating very few possible regions where *Arabis auriculata* could survive the LGM; (C) Modelled current distribution of suitable niches for *Arabis montbretiana* and (D) distribution of suitable niches during the LGM; the modelled distribution clearly indicates that it is restricted to Asia minor and reaches Eastwards into Central Asia (E) Modelled current distribution of suitable niches for *Arabis iberica* and (F) distribution of suitable niches during the LGM; suitable niches nowadays would also occur in North Africa and even reach the Balcans; however, the taxon does not occur there (G) Modelled current distribution of suitable niches for *Arabis nova* and (H) distribution of suitable niches during the LGM;the present distribution of suitable niches is very narrow and mosty not overlapping with the suitable niches for *Arabis iberica*

In summary we could identify mostly non-overlapping species distribution ranges based on recent bioclimatic variables as well as putative areas where our annual species in this study species could survive the LGM which fit with known refuge areas.

### Taxonomic differentiation of Arabis iberica and Arabis nova and a new combination

*Arabis iberica* has long been treated as a subspecies of *Arabis nova*. Past (Karl et al. 2012; Karl and Koch 2013) studies have already shown that *Arabis nova* is paraphyletic with *Arabis nova* ssp. *nova* being closely related to *Arabis auriculata* while *Arabis nova* ssp. *iberica* forms a clade with *Arabis montbretiana* and *Arabis kennedyae*. In our study we have confirmed this finding (Supplementary Figure 1) and that the groups containing *Arabis nova* and *Arabis iberica* diverged already ∼11 mya (Figure 1). Further our niche modelling indicates that the most likely distribution ranges determined by niche modelling of both taxa do only overlap in a small region in Northern Spain. Based on these findings we raise *Arabis nova* ssp. *iberica* to species rank as *Arabis iberica* (Rivas Mart. Ex. Talavera)

M. Koch, R. Karl, D. German et Al-Shehbaz:

*Arabis iberica* (Rivas Mart. ex Talavera) M. Koch, R. Karl, D. German et Al-Shehbaz, comb. et stat. nov.

*≡ Arabis nova* subsp. *iberica* Rivas Mart. ex Talavera in **Anales Jard. Bot. Madrid 50: 148 (1992)\***

*≡ Arabis nova* subsp. *iberica* Rivas Martínez in **Anales Inst. Bot. Cavanilles 21: 221 (1964)**, nom. inval.

*Holotypus:* “Flora Hispánica - Herbario Normal / Centuria VII Abril de 1951 / 631. *Arabis nova* Vill. var. *serrifera* F.Q. / Prov. de Ciudad Real: La Calderina, schistous slopes, at 1,100 m.s.l. / recorded and collected by F.Q. on May 31rd 1924 / cultivated in Barcelona in April 1925.” (MA 174931; isotype – SANT 6605).

*≡ Arabis nova* Vill. var. *serrifera* F.Q., nomen nudum

= *Arabis saxatilis f. latiorifolia* Pau, nomen nudum (see Nualart et al. 2021:14) in **Iter Marocc.: 198 1930 (1929)**, Herbarium G, G00424997, leg. P. Font Quer no. 198, Morocco 06.06.1929 (Maroc, hab. in cedretis montis Tidiguin, 2000 a.s.l.

The native range of the taxon is Central & S. Central Spain, Morocco. It is an annual or biennial and grows primarily in the temperate biome.

#### TYPE SPECIMENS (Supplementary File 5)

Madrid (MA):

https://imagenes.rjb.csic.es/herbarioV/visorVCat.php?img=MA-01-00174931 (holotype) https://imagenes.rjb.csic.es/herbarioV/visorVCat.php?img=MA-01-00340667 (paratype)

Santiago (Sant): as *Arabis nova* Vill. var *serrifera* F.Q. https://plants.jstor.org/stable/viewer/10.5555/al.ap.specimen.sant6605?page=1 (isotype) https://plants.jstor.org/stable/viewer/10.5555/al.ap.specimen.sant6606?loggedin=true (paratype)

Barcelona (BCN): as *Arabis saxatilis* All. f. *latiorifolia* Pau (erroneously labeled as syntype, nomen nudum)

https://plants.jstor.org/stable/viewer/10.5555/al.ap.specimen.bcn72336 (syntype)

Geneve (G): as *Arabis saxatilis* All. f. *latiorifolia* Pau (erroneously labeled as holotype, nomen nudum)

https://www.ville-ge.ch/musinfo/bd/cjb/chg/pics.php?lang=en&id=G00424997&iip=cjbiip/cjb44/img_212/G0042499 7.ptif

Nualart N, Soriano I, Pérez Orieto D, Ibánez N (2021) Cataloque and typification of the Moroccan taxa published by Carlos Pau. Phytotaxa 519 (1: 1-94.)

*[Salvador Talavera: Notulae Taxinomicae, Chorologicae, Nomenclaturales, Bibliographicae aut Phologicae in Opus “Flora Iberica” Intendentis]

## Discussion

Due to its simple genetic basis which was revealed recently (Zhai et al. 2024) switches between annual and perennial life history are frequent in the Brassicaceae. Although annuals are rare in the tribe Arabideae when compared to the rest of the family (8% vs. 25%) switches have occurred multiple times independently. Two of the annual lineages are closely related to the perennial model species *Arabis alpina*. So far representatives of the four taxa making up these two lineages had been subject to comparative molecular analyses. However, not much was known about the evolutionary history of these lineages. Therefore, inhere we focused on shedding some light on the past of *Arabis auricluata*, *A. iberica*, *A. montbretiana* and *A. kennedyae*.

Past climatic oscillations are a major determinant of nowaday’s species distribution. The Quaternary ice ages forced plant and animal species into glacial refugia which in Europe were typically located in the Mediterranean or on the Iberian Peninsula (Avise, 2000; Hewitt, 2004). However, refuge areas were also identified along the margin of the Alps (Schönswetter et al. 2005) and so were events of in situ survival (e.g. Parsiod and Besnard 2007).

Previously we had analysed the phylogeographic history of Arabi alpina, a model perennial from the genus (Koch et al. 2006). We had identified its origin in Asia minor and detected three migration waves of which two lead to the Arabian penisula and East Africa while the third one lead into entire Europe, Iceland and also Greenland. Glacial refugia were hepothesized to have been located in the Mediterranean. Later studies identified Italy as a major long term refuge area (Ansell et al. 2008). Thereby *Arabis alpina* presents a typical example for the phylogeographic history of a perennial plant taxon affected by the Quaternary climatic oscillations.

In the current study we were interested in looking at the phylogeography in *Arabis* from an annual perspective. Therefore, we sampled the two annual lineages which are in fact both closely related to *Arabis alpina* (Karl and Koch 2013). However, both lineages show very distinct distribution patterns, suggesing that their phyogeographic history might differ significantly as well.

The first lineage we focused on comprised only *Arabis auriculata*, an annual with a broad Eurasian distribution. *Arabis auriculata* split from its annual sister *Arabis nova* ssp. *nova* about 9 mya and started diversifying around 1.2 mya, a period referred to as Mid-Pleistocene Transition which was characterized by a gradual increase in length of glaciation cycles (Clark et al. 2006).

Our findings indicate that *Arabis auriculata* exhibits a continuous distribution pattern, with a geographically structured haplotype distribution. Three major haplotypes were observed in *Arabis auriculata*: one exclusive to Asia Minor, one spanning southern and central Europe, and another confined to the Iberian Peninsula along with its derived singletons. These results suggest historical processes of regional diversification and maybe limited long-distance dispersal in this species.

The highest haplotype diversity was detected on the Iberian Peninsula and in North Africa, where numerous singleton haplotypes were identified. A high genetic diversity on the Iberian Peninsula has been identified for multiple taxa e.g. for Arabidopsis thaliana (Pico et al. 2008), Artemisia crithmifolia (García-Fernández et al. 2017), *Arabis scabra* (Koch et al. 2020) or Juniperus phoenicea (Dzialuk et al. 2011). For the former three the Iberian Peninsula is assumed to have acted as glacial refuge area which lead to the preservation of a higher genetic diversity while for the latter taxon the high genetic diversity is explained by the complex topography of the region and its long-term stable climatic conditions. In the case of *Arabis auriculata* it may have been a mixture of both, topographic diversity and the Iberian Peninsula acting as glacial refuge area. The flora of the Iberian Peninsula shows a strong connection to the flora of North Western Africa (Molina-Venegas et al. 2015). Geographical structuring of populations seems to be strong in North Western Africa and was detected for example in Arabidopsis thaliana (Brennan et al. 2014). In this case also a strong genetic connection was detected between the Iberian Peninsula or more general the Western Mediterranean and Morocco. However, in the case of A. auriculata our phylogenetic analyses suggest that the lineage rather arrived from the Eastern Mediterranean than from the Iberian Peninsula.

The second lineage we analysed comprised *Arabis montbretiana*, *Arabis kennedyae* and *Arabis iberica*, three annuals with distinct and - compared to *Arabis auriculata* - narrow distribution ranges. If the lineage is considered as a whole it shows the hallmarks of an Iberian-Euxinian disjunction like in *Rhododendron ponticum* ssp. *baeticum* (Mejias et al. 2007). In our case it means that the part of the lineage comprising *Arabis iberica* is confined to the Iberian Peninsula while the part of the lineage comprising *Arabis montbretiana* and *Arabis kennedyae* is found in the Eastern Mediterranean (and here into Central Asia as well). The Iberian-Euxinian disjunction is a result of the cooling climate that started 33.5 mya and then continued (Leutert et al. 2020). It lead to the extinction of the typical vegetation of the Tertiary in most parts of Europe. Our divergence time analysis suggests that the lineage comrprising the three annual taxa mentioned before split about 1.7 mya. So the disjunction happened in the Quaternary which means that also events of that period may lead to Iberian-Euxinian disjunctions.

*Arabis iberica* has formed three haplotypes on the Iberian Peninsula where haplotype Ai Hap A is the most frequent haplotype and Ai Hap B and Ai Hap H are less frequent or a singleton. The split between Ai Hap H which is found in Northern Spain and the two other haplotypes which are found in Central Spain seems to have happened already 1.3 mya. This suggests that (a) migration into the Iberian Peninsula happened rather quickly as the split between *A. iberica* on the one hand and *A. montbretiana/kennedyae* on the other hand happened 1.7 mya and (b) that in the Quaternary two *A. iberica* lineages were already present on the Iberian Peninsula, one in Northern Spain, the other one in Central Spain. Ai Hap B split from Ai hap A about 18,000 years ago, which is post glacial and may correspond to a postglacial migration event when taxa could leave their refuge areas again. The split between Central Iberian and North Iberian accessions is not uncommon and was detected also in Arabidopsis thaliana (Pico et al. 2008), however the genetic cluster detected in the North of the Iberian Peninsula was also found in the South but not in the Center.

The haplotype patterns in *A. montbretiana* suggest a gradual eastward migration from Asia Minor into Central Asia. The split between *A. iberica* and *A. montbretiana* happened 1.7 mya, the split of Am Hap D (Arabian Peninsula and Iran) and Hap C (coast of Turkey and Cyprus) happened 1.6 mya. Further splits of haplotypes could not be dated. Migrations of taxa from the Eastern Mediterranean into Central Asia were common during the Pleistocene. The Irano-Turanian region may have acted as a significant corridor for herbaceous plant migration, facilitating the movement of species from Central Asia and Afghanistan into the Iranian plateau and vice versa and further into the (Sub)Mediterranean, as well as contributing to the overall high levels of plant endemism and diversity within these region. Its geoclimatic conditions and mountain systems create diverse habitats and act as both refuges and barriers, shaping the distribution patterns of its native and naturalized vascular plants, which include therophytes (annual plants) and hemicryptophytes (herbaceous perennials) (e.g., Memariani et al. 2016, Manafzadeh et al. 2014).

Am Hap C is shared by *A. montbretiana* and *A. kennedyae* which is endemic to Cyprus. Whole plastome data indicate that *A. montbretiana* and *A. kennedyae* actually split 157 kya. This time coincides with the Penultimate Glacial Maximum during which sea levels in the Mediterranen were considerably lowere therefore decreasing the distance between mainland and Cyprus (Menviel et al. 2019). However, Cyprus has not been connected to the main land since at least 5.3 my (Constantinou and Panayides 2013) but still lower sea levels may have facilitated an easier long distance dispersal from the mainland to Cyprus.

Our ecological niche modeling further supports these findings by demonstrating the varying habitat availability for the studied species. The results indicate that suitable habitats for *Arabis auriculata* extend across Europe and Central Asia, mirroring its broad present-day distribution. In contrast, the predicted suitable habitats for *A. montbretiana* and *A. iberica* are largely restricted to Asia Minor and Central Asia or the Iberian Peninsula and Northwest Africa, respectively. This pattern suggests that environmental constraints may have played a key role in the divergence and limited dispersal of these lineages, potentially leading to their present-day geographical separation.

Overall, our study highlights the importance of combining phylogenomic data, haplotype network analyses, and ecological niche modeling to reconstruct the evolutionary history of plant lineages. The observed patterns of genetic divergence and ecological constraints in *A. montbretiana* and *A. iberica* reinforce the role of biogeographic barriers in shaping species distributions. Future studies incorporating broader taxonomic sampling and additional environmental variables could further refine our understanding of the evolutionary processes driving diversification in the tribe Arabideae.

## Acknowlegments

We acknowledge financial support from German Research Foundation (DFG) with grant no. KO2302/23-2.

## Copyright

The copyright holder for this preprint is the author/funder, who has granted bioRxiv a license to display the preprint in perpetuity. All rights reserved. No reuse allowed without permission.

## Supplementary Materials

**Supplementary Figure 1** Maximum likelihood phylogenetic reconstruction based on coding sequences from plastid genomes. Bootstrap support values are colour coded (legend in figure); the lineages recognized are in agreement with previous studies (e.g. Karl and Koch 2013).

**Supplementary Figure 2** Maximum likelihood tree based on the *trnLF region* including data from all plastomes generated within this study and additional data from previosu studies (Karl and Koch 2013; Koch et al. 20016).

**Supplementary Figure 3** Maximum likelihood tree based on the *ITS* region including all accessions for which sequencing data were generated within this study; the tree is used as a barcoding phylogeny and for hybrid identification by comparison to the tree based on the *trn*LF region.

**Supplementary Figure 04** Correlation plot for identifying correlated bioclim variables.

**Supplementary File 1** Alignment of all coding sequences extracted from the plastomes; the alignment was used for maximum likelihood and BEAST analysis. Partitions are given in Supplementary File 2.

**Supplementary File 2** Partitions corresponding to the alignment in supplementary File 1.

**Supplementary File 3** Alignment of the ITS region corresponding to the maximum likelihood analysis shown in supplementary Figure 3.

**Supplementary File 4** Alignment of the *trn*LF region corresponding to the maximum likelihood analysis shown in supplementary Figure 2.

**Supplementary File 5** Images of *A. iberica* type material vouchers.

**Supplementary Table 1** Information on all accessions used within this study (HEID herbarium code, Genbank accession codes, information linked to material collection, read numbers (where applicable).

**Supplementary Table 2** Calibration Points used in the BEAST analysis as extracted from the BEAST analysis in Hohmann et al. (2015).

