## Supplementary figures and images for "Split or spread - A spatio-temporal framework for the evolution of annual *Arabis* (Brassicaceae) in Eurasia"

### Supplementary Figure 1

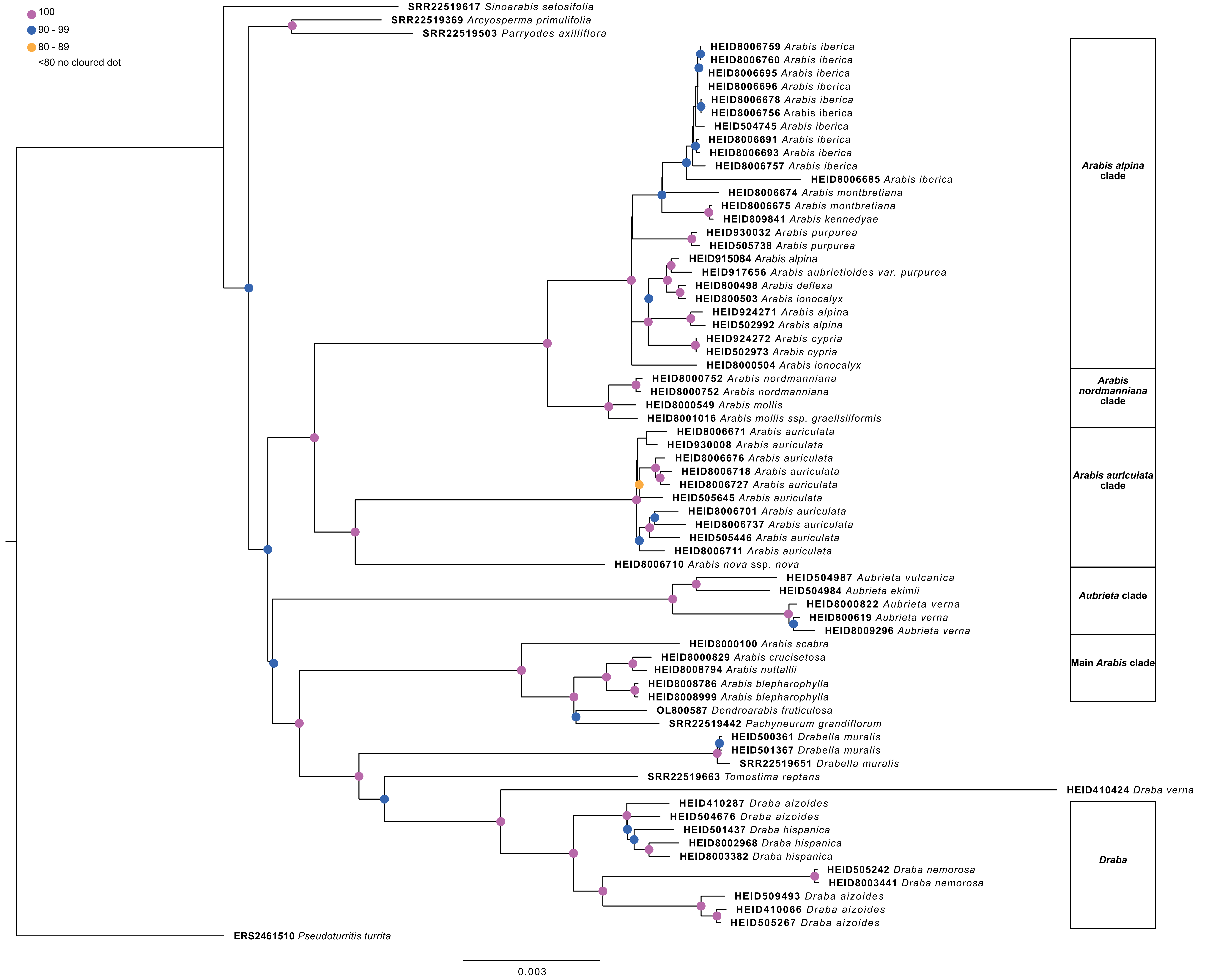

### Supplementary Figure 2

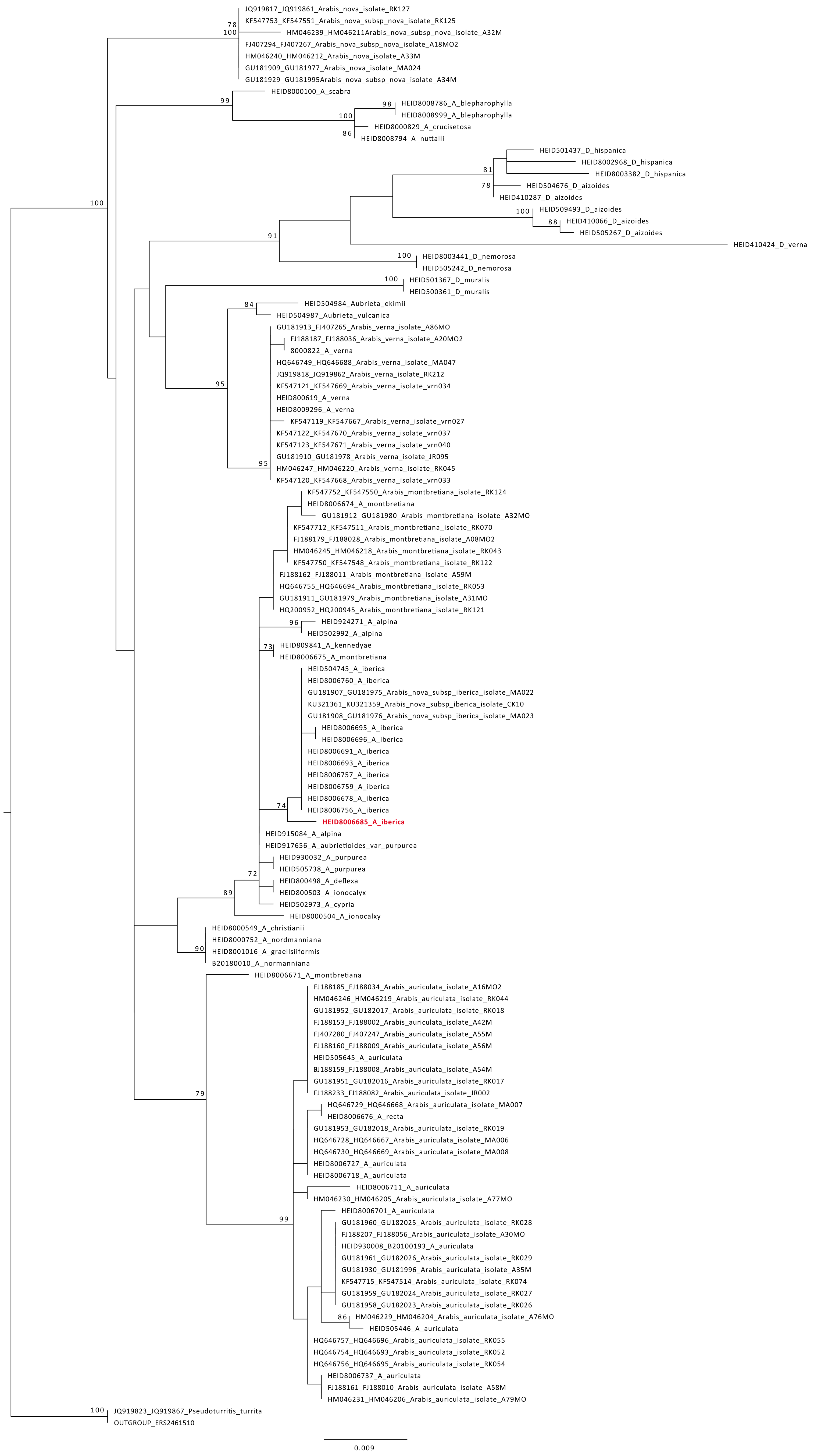

### Supplementary Figure 3

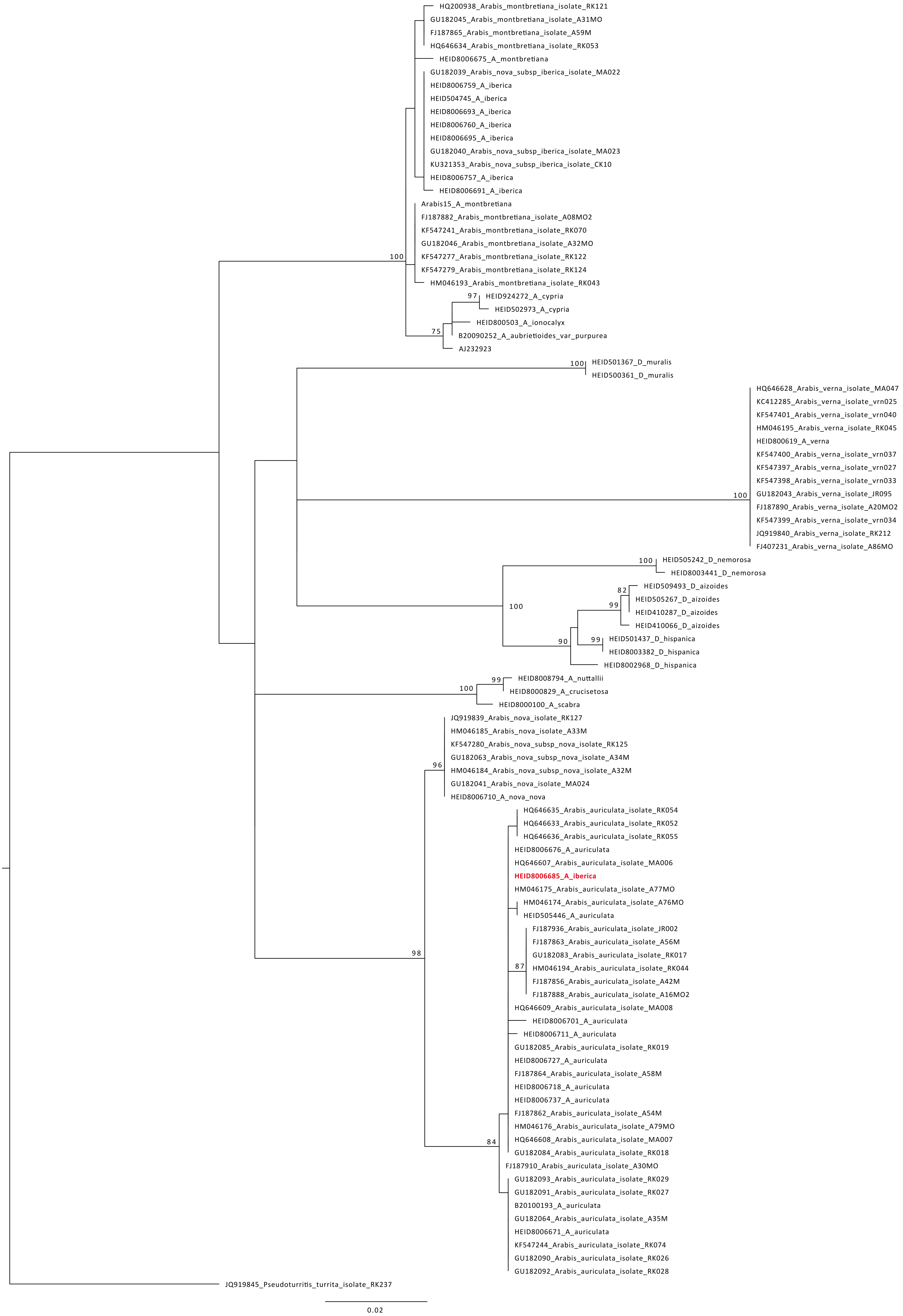

### Supplementary Figure 4

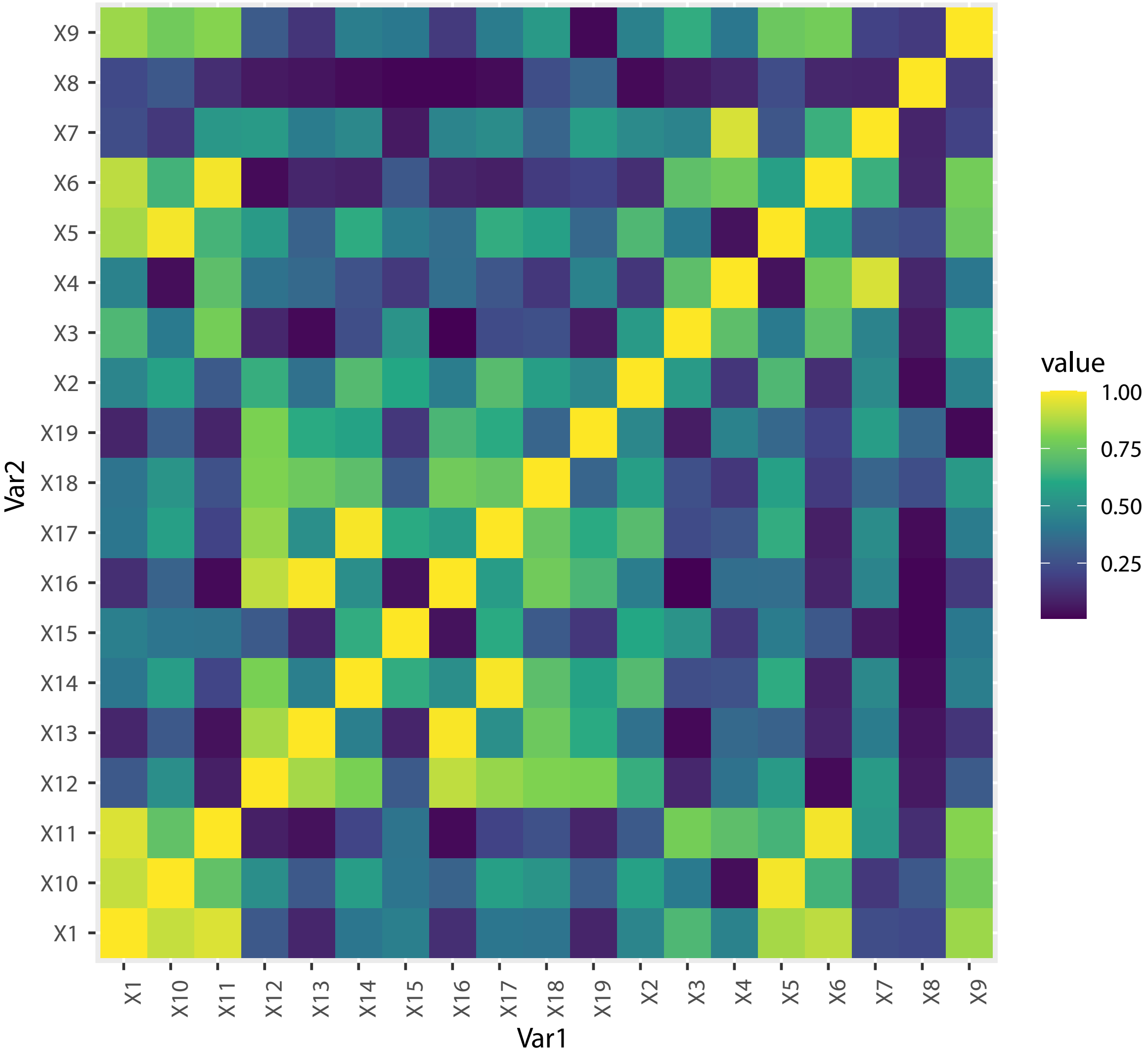
